# Specific connectivity optimizes learning in thalamocortical loops

**DOI:** 10.1101/2022.09.27.509618

**Authors:** Kaushik Lakshminarasimhan, Marjorie Xie, Jeremy Cohen, Britton Sauerbrei, Adam Hantman, Ashok Litwin-Kumar, Sean Escola

## Abstract

Cortico-thalamo-cortical loops have a central role in cognition and motor control, but precisely how thalamus contributes to these processes is unclear. Recent studies showing evidence of plasticity in thalamocortical synapses indicate a role for thalamus in shaping cortical dynamics – and thus behavior – through learning. Since corticothalamic projections compress cortical activity into a lower-dimensional thalamic activity space, we hypothesized that the computational role of thalamus would depend critically on the structure of corticothalamic connectivity. To test this, we identified the optimal corticothalamic structure that promotes biologically plausible learning in thalamocortical synapses. We found that corticothalamic structures specialized to carry an efference copy of the cortical output benefit motor control, while corticothalamic connections that communicate the directions of highest variance in cortical activity are optimal for working memory tasks. We analyzed neural recordings from mice performing grasping and delayed discrimination, and found corticothalamic interactions consistent with these predictions. These results suggest that thalamus orchestrates cortical dynamics in a functionally precise manner through structured connectivity.

## 2 Introduction

Recent years have seen a renewal of interest in understanding the role of higher-order thalamus in mammalian behavior. Unlike primary thalamic nuclei, such as the lateral geniculate nucleus, which relays information from the sensory periphery to cortex, higher-order nuclei do not receive inputs from the periphery. Consequently, the conventional view of thalamus as a relay has been revised to accommodate a role for higher-order thalamus as a “higher-order relay” that transmits information from one cortical area to another (Sherman, 2007; Sherman & Guillery, 2018). Meanwhile, molecular tracing studies have revealed a frequent motif of reciprocal interactions in which cortical projections to the thalamus target nuclei that project back to the same cortical area. Such reciprocal cortico-thalamo-cortical (CTC) loops have been observed in rodent sensory (Guo, Yamawaki, Barrett, Tapies, & Shepherd, 2020), motor (Yamawaki & Shepherd, 2015; Guo, Yamawaki, Svoboda, & Shepherd, 2018; Economo et al., 2018; Muñ oz-Casta ñeda et al., 2021), and prefrontal (Collins, Anastasiades, Marlin, & Carter, 2018) cortices and stand in contrast to a view of such nuclei as higher-order relays across cortical areas. Recent physiological studies have demonstrated that these loops are required for many cognitive functions (Bolkan et al., 2017; Guo et al., 2017; Schmitt et al., 2017; Rikhye, Gilra, & Halassa, 2018; Alcaraz et al., 2018). Furthermore, the evolution of cortical activity depends on thalamus (Sauerbrei et al., 2020) and inhibiting higher-order thalamus suppresses cortical activity (Guo et al., 2017; Schmitt et al., 2017). However, the specific computational role of cortico-thalamo-cortical loops is not fully understood.

Reciprocal CTC loops represent a departure from the labelled line view of thalamus as they feature signal compression (from cortex to thalamus) and expansion (from thalamus to cortex). Normative models developed to clarify the role of compression and expansion in feedforward neural networks have improved our understanding of the computations in many brain areas including the retina (Zhaoping, 2006; Druckmann, Hu, & Chklovskii, 2012), primary visual cortex (Olshausen & Field, 1996; Zhu & Rozell, 2013), olfactory bulb (Zhang & Sharpee, 2016; Qin, Li, Tang, & Tu, 2019) and the cerebellum (Litwin-Kumar, Harris, Axel, Sompolinsky, & Abbott, 2017; Muscinelli, Wagner, & Litwin-Kumar, 2022; Xie, Muscinelli, Harris, & Litwin-Kumar, 2022). For instance, theories of compressed sensing and efficient coding have shown that whereas random compression can preserve the similarity structure of sparse representations, the optimal compression strategy is to extract the principal components when inputs are strongly correlated (Ganguli & Sompolinsky, 2012). Unfortunately, insights gained from analyzing feedforward networks cannot be directly applied to understand signal transformation in CTC loops due to their recurrent processing.

We and others have shown in prior theoretical studies that it is possible to perform computations by tuning synaptic weights in the CTC loop appropriately (Kao, Sadabadi, & Hennequin, 2021; Logiaco, Abbott, & Escola, 2021), consistent with experimental studies showing that thalamocortical synapses exhibit plasticity in many areas of the adult brain (Herry, Vouimba, & Garcia, 1999; Oberlaender, Ramirez, & Bruno, 2012; Biane, Takashima, Scanziani, Conner, & Tuszynski, 2016; Audette, Bernhard, Ray, Stewart, & Barth, 2019; Hasegawa, Ebina, Tanaka, Kobayashi, & Matsuzaki, 2020; Adam, Schaeffer, Johnston, Menon, & Everling, 2021; Sohn et al., 2022). Such modifications can be viewed as low-rank weight modifications to the connectivity of cortical recurrent neural networks (Mastrogiuseppe & Ostojic, 2018; Schuessler, Mastrogiuseppe, Dubreuil, Ostojic, & Barak, 2020; Dubreuil, Valente, Beiran, Mastrogiuseppe, & Ostojic, 2022). However, these models have not addressed how these synapses may be updated with biologically plausible learning rules that operate using locally available signals. In particular, we show that standard approaches for local, biologically plausible learning in recurrent neural networks fail to optimize corticothalamic connectivity, because accurate credit assignment at these synapses requires knowledge of the global structure of network activity.

In this study, we hypothesize that the ability of the thalamus to contribute to learning depends critically on the structure of corticothalamic synapses and ask what forms of corticothalamic connectivity structure optimize learning. We use a recently proposed biologically plausible, three-factor supervised learning rule (Murray, 2019) to update thalamocortical synapses, while assuming that corticothalamic synapses are optimized on a slower evolutionary or developmental timescale. We separately train models to perform two different tasks – autonomous motor control and working memory – and determine the optimal corticothalamic structure in each case (Wang, 2021; Hospedales, Antoniou, Micaelli, & Storkey, 2022). We find that the optimal corticothalamic connectivity is structured and task-dependent. Specifically, learning motor control is optimized by corticothalamic connections specialized to carry an efferent copy of the muscle command, while learning to perform working memory is optimized by corticothalamic connections that convey modes of cortical activity with the highest variance. We analyzed neural recordings from mice performing grasping and delayed discrimination tasks (Guo et al., 2017; Sauerbrei et al., 2020), and found that the influence of cortical activity on thalamus is consistent with this model. These results suggest that thalamus orchestrates cortical dynamics in a functionally precise manner through structured connectivity.

## 3 Results

We begin by constructing a model combining cortex and thalamus based on two fundamental properties of thalamic nuclei that distinguish them from the cortex. First, there are far fewer neurons in the thalamus (Halley & Krubitzer, 2019) which acts as a structural bottleneck in this loop. Second, local recurrent excitation is a defining feature of the cortex but absent within the thalamus (Arcelli, Frassoni, Regondi, Biasi, & Spreafico, 1997; Halassa & Sherman, 2019). Therefore, we consider a model in which a network of *N* interconnected cortical neurons is reciprocally connected with *M* uncoupled thalamic neurons, with *M* ≪ *N* (**Figure 1A**). The cortical population activity is denoted by **h** *t*∈ ℝ ^*N*^, thalamic population activity by **r** *t* ∈ ℝ^*M*^, and *S* external inputs (if any) by **x** *t*∈ ℝ^*S*^. The cortical activity evolves according to:

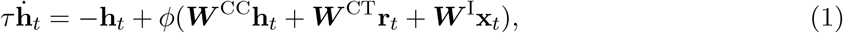

where subscripts denote time, *τ* is the neuronal time constant, and the nonlinearity *ϕ*(·) is the tanh function. ***W*** ^CC^ ∈ ℝ ^*N* ×*N*^, ***W*** CT ∈ ℝ ^*N* ×*M*^ and ***W*** ^I^ ∈ ℝ ^*N* ×*S*^ denote corticocortical, thalamocortical and input weight matrices respectively. Thalamic activity, on the other hand, is assumed to depend only on cortical activity as **r** *t* = *ϕ*(***W*** ^TC^ **h** *t*) where ***W*** ^TC^ ∈ ℝ ^*M* ×*N*^ 1denotes the matrix of corticothalamic weights. Therefore, the thalamic membrane potential **v** *t* = ***W*** ^TC^**h** _*t*_is an *M* - dimensional projection of the cortical activity. Network output **y** _*t*_∈ ℝ^*R*^, which we optimize to perform behavioral tasks, is modeled as a linear readout of the cortical activity, **y** _*t*_ = ***W*** °**h** _*t*_with readout weights ***W*** °∈ ℝ^*R*×*N*.^

**Figure 1:**
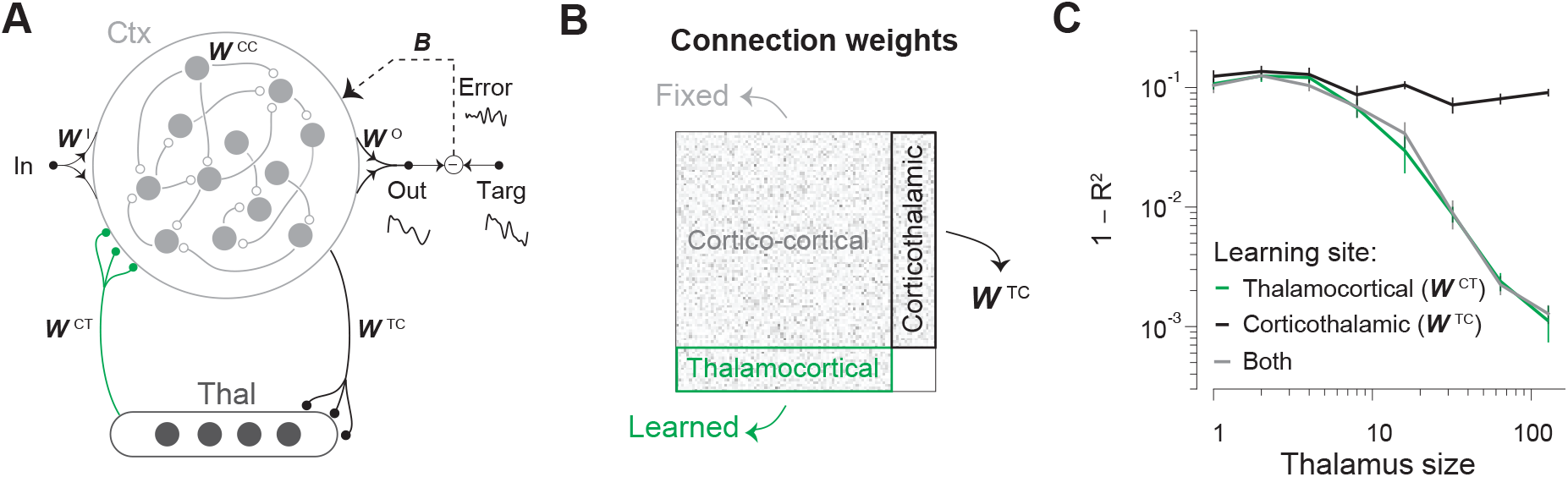
Combined model of cortex and thalamus. **A**. Schematic illustration of a thalamocortical network model. **B**. Synaptic weights of the model. Corticocortical weights (***W*** ^CC^) are fixed. Local learning rules can be used to update thalamocortical weights (***W*** ^CT^), but it is not known whether such rules are effective for corticothalamic weights (***W*** ^TC^). **C**. Learning performance as a function of thalamic population size when learning is mediated by a local plasticity rule (RFLO) in corticothalamic weights (black), thalamocortical weights (green), or both (gray). Models have *N* = 256 cortical neurons. The number of thalamic neurons (*M*) is varied from 1 to 128. Error bars denote standard errors estimated by bootstrapping.

In this model, cortical neurons interact directly via corticocortical connections and indirectly via the thalamic bottleneck, which compresses the cortical signal into an *M* -dimensional space (through ***W*** ^TC^) before expanding it back into an *N* -dimensional space (through ***W*** ^CT^). The effective interaction weights can be expressed as a sum of recurrent connectivity within the cortex and weights in the CTC loop, ***W***_eff_ = ***W*** ^CC^+ ***W*** ^CT^***V W*** ^TC^where ***V*** = diag(*ϕ*^′^(**v** _*t*_)) is an *M* × *M* diagonal matrix composed of the derivative of the activity of thalamic neurons. This effective interaction matrix can be viewed as a low-rank (rank-*M*) perturbation of the *N* × *N* cortical connectivity matrix ***W*** ^CC^. Such models are interesting from a computational perspective because recent theoretical work has shown that many tasks can be solved by combining a full-rank random component with appropriate low-rank connectivity structures (Mastrogiuseppe & Ostojic, 2018) and that gradient-based learning typically induces low-rank changes in recurrent weights (Schuessler, Mastrogiuseppe, et al., 2020). We ask whether low-rank CTC connectivity can be learned to support behavioral tasks in a biologically plausible manner within our model.

### 3.1 Thalamocortical learning rule

Recent experimental evidence suggests that thalamocortical plasticity continues into adulthood and is a major substrate for learning (Oberlaender et al., 2012; Biane et al., 2016; Audette et al., 2019; Hasegawa et al., 2020; Adam et al., 2021; Sohn et al., 2022), and inhibiting thalamus impairs learning (Tanaka et al., 2018). Accordingly, we assume that thalamocortical synapses (***W*** ^CT^, **Figure 1B** – green) are adjusted such that the overall model minimizes Σ _*t*_ ∥***ε*** _*t*_ ∥ ^2^, where 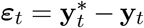 denotes the mismatch between the network output **y** _*t*_ and some desired target output function 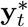. Specifically, we train thalamocortical synapses in the model using a local, biologically plausible learning rule based on a recently proposed algorithm (Random-Feedback-Local-Online, or RFLO, learning; see Methods) (Murray, 2019).

The synaptic weight updates in this scheme are derived by making two approximations to algorithms that minimize squared-error by computing the full gradient with respect to the weights (Williams & Zipser, 1989). The first approximation consists of dropping nonlocal terms from the gradient, so that computing the update to a given thalamocortical synapse requires only presynaptic (thalamic neuron) and postsynaptic (cortical neuron) activities, rather than information about the entire state of the cortex including all of its synaptic weights. The second is to project the error ***ε*** _*t*_ back into the cortical network for learning using a random feedback pathway with weights ***B*** ∈ ℝ ^*N* ×*R*^, rather than feedback weights that are precisely tuned to match the readout weights (i.e. ***B***^*T*^ ≠ ***W*** °). This relaxation is made possible by learning readout weights ***W*** ° in conjunction with thalamocortical weights, such that alignment between readout and feedback weights can be established during the course of learning (Lillicrap, Cownden, Tweed, & Akerman, 2016). The resulting update rule for thalamocortical weights is given by:

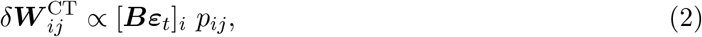

where 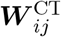 denotes the synaptic weight from thalamic unit *j* onto cortical unit *i*. The eligibility trace *p_*ij*_ reflects the correlation between recent activity of the thalamic unit *j* and the cortical unit 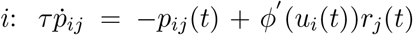. Readout weight updates follow a standard delta-rule, 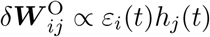 where *ε* _*i*_ (*t*) denotes the error in the *i*^th^*output dimension. Like other three-factor rules (Gerstner, Lehmann, Liakoni, Corneil, & Brea, 2018), the update rule for thalamocortical synapses depends only on the presynaptic activity (thalamic neuron), the postsynaptic activity (cortical neuron), and the error signal at each moment (Methods). Evidence for error signals in the superficial layers of the cortex (Inoue, Uchimura, & Kitazawa, 2016; Heindorf, Arber, & Keller, 2018) suggests that thalamocortical synapses located on apical dendrites e.g. (Guo et al., 2018) are particularly good candidates for this learning rule, but apical error signals could also drive plasticity in basal thalamocortical synapses via dendritic plateau potentials (Guerguiev, Lillicrap, & Richards, 2017; Bittner, Milstein, Grienberger, Romani, & Magee, 2017).

We found that the performance of models trained by applying the above learning rule improves with the number of thalamic neurons, suggesting that local plasticity at thalamocortical synapses (***W*** ^CT^) faciliates learning (**Figure 1C** – green). This improvement in learning performance is accompanied by an increase in the alignment between weights in the feedback pathway **B** (which communicate error signals to post-synaptic neurons) and readout weights **W**°(Figure S1B – green), indicating that the learning rule performs credit assignment – conveying appropriate learning signals to neurons upstream of behavioral output. Nonetheless, the performance of this learning strategy may depend critically on the signal received by thalamic neurons via corticothalamic projections (***W*** ^TC^, **Figure 1B** – black). Studies of biologically plausible algorithms in recurrent neural networks have typically dealt only with the learning of synapses onto neurons that are directly connected to the readout (Miconi, 2017; Alemi, DenÈve, Machens, & Slotine, 2018; Murray, 2019; Gilra & Gerstner, 2017). Since corticothalamic synapses are further upstream than thalamocortical synapses, it is not known whether biologically plausible plasticity rules can perform credit assignment in these synapses. To test this, we trained our model by applying a local plasticity rule analogous to Equation 2 to update corticothalamic synapses (Methods – Equation 6, Figure S1A). In contrast to models that learn by updating thalamocortical synapses, the resulting model fails to learn even when the thalamic population size is comparable to the size of the cortex (**Figure 1C** – black). In fact, local plasticity operating simultaneously at both thalamocortical and corticothalamic synapses does not yield any improvement in learning performance over what is obtained from learning only thalamocortical synapses with fixed random corticothalamic connections (**Figure 1C** – gray vs green).

The failure of local plasticity in corticothalamic synapses to improve performance can be understood by comparing the feedback alignment of neurons in cortex and thalamus. Whereas local plasticity achieves a high level of feedback alignment in cortical neurons, the alignment is no greater than chance in thalamic neurons (Figure S1B). Therefore, error signals cannot mediate learning via local plasticity in corticothalamic synapses (Figure S1C). In general, performing accurate credit assignment for signals in brain areas that do not directly drive behavior requires accounting for the downstream transformations performed on these signals prior to their influence on behavior. In our model, this means that weight updates in corticothalamic synapses would need to be guided by error signals gated through both the time-varying neural activity in the cortex and thalamocortical weights, a requirement that cannot be easily implemented in a biologically plausible way.

### 3.2 Optimal corticothalamic connectivity

Since biologically plausible feedback-driven plasticity was unable to optimize corticothalamic connectivity, we asked whether other mechanisms could select patterns of synaptic weights that improve performance. In particular, we sought to identify subspaces of cortical activity that, when communicated to the thalamus, improve learning supported by thalamocortical plasticity (**Figure 2A**). Specific corticothalamic connectivity that routes activity in such subspaces may be established by evolutionary or developmental processes, or by plasticity rules of a different form than Equation (2).

**Figure 2:**
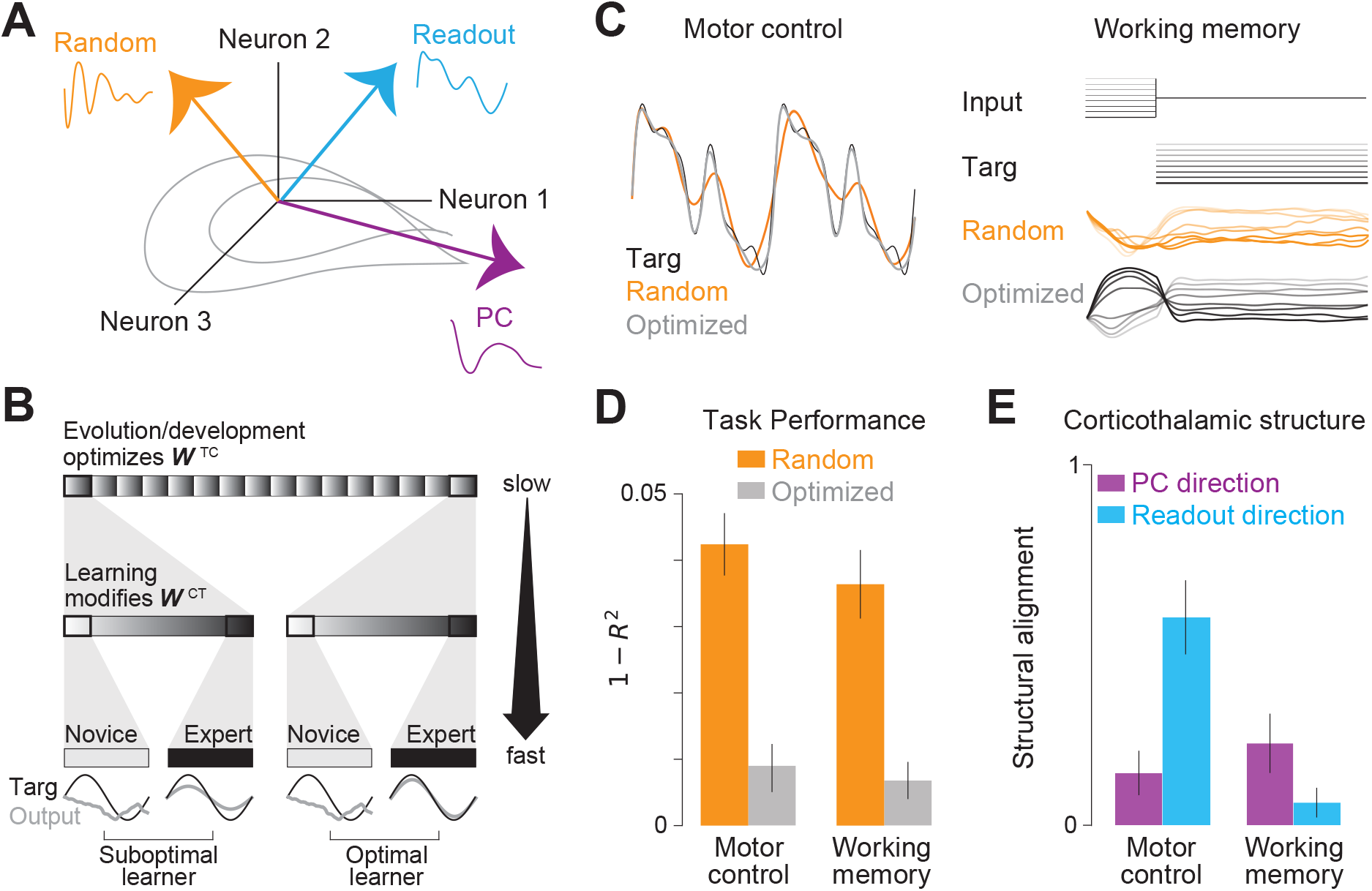
Corticothalamic connectivity optimized by meta-learning. **A**. Corticothalamic weights may prioritize communicating specific subspaces of cortical activity, such as the readout (cyan) or principal component directions (purple), in contrast to a random subspace (orange). **B**. Schematic of meta-learning procedure. Optimizing corticothalamic weights on a slow timescale (top, “outer loop”) improves error-driven learning in thalamocortical synapses (middle, “inner loop”), improving performance (bottom). **C**. Left: Median outputs (across simulations) of networks trained using meta-learned corticothalamic weights (gray) and random corticothalamic weights (orange), on the autonomous control task. Black line denotes the target function. Right: Inputs and outputs of networks trained using meta-learned corticothalamic weights (gray) and random corticothalamic weights (orange) for the different conditions of the working memory task. **D**. Task average performance error of the model (across simulations initialized with different thalamocortical weights) trained using the meta-learned corticothalamic weights (gray), compared to the average performance error of the model trained using random corticothalamic weights (orange). **E** Alignment of meta-learned corticothalamic weights with the readout direction (cyan) and the direction of the leading principal component of cortical activity (purple) in the two tasks. Error bars denote ± 1 SEM estimated by bootstrapping. All models have *N* = 256 cortical neurons and *M* = 32 thalamic neurons.

To begin, we took an approach known as meta-learning or “learning to learn” (Wang, 2021; Hospedales et al., 2022). Corticothalamic weights were optimized at a longer time scale across thousands of epochs (“outer loop”), each of which comprises a few hundred trials of thalamocortical learning (“inner loop”) (**Figure 2B**; Methods). The outer loop represents the evolutionary or developmental processes that optimize corticothalamic connectivity. Optimization was performed via backpropagation through time. To ensure that corticothalamic weights discovered by this technique did not depend on the precise state of thalamocortical weights at the beginning of learning, we reset the thalamocortical weights at the beginning of each epoch. Moreover, to ensure that this technique promotes learning specifically via thalamocortical synapses, we fixed all other weights, and enforce feedback alignment (***B***^T^= ***W*** °) during the meta-learning procedure. We later relax these constraints and test our conclusions in a model with plasticity in corticocortical and readout weights. To test the learning performance, we train the thalamocortical weights in the model using the fixed, optimized corticothalamic weights and compare the performance against a model trained using fixed random corticothalamic weights (Methods).

Which corticothalamic projections are most suitable for thalamocortical learning could depend on the task. To test this hypothesis, we consider two prototypical neuroscience tasks with distinct computations – an autonomous motor control task and a working memory task. The goal of the autonomous control task is to output a complex temporal waveform without the aid of external inputs (i.e., analogous to the EMG activity required for internally generated movement). The goal of the working memory task is to output the amplitude of one of eight possible transient input pulses during a subsequent delay period (Methods). We optimized corticothalamic weights for each task as outlined above, and found that meta-learned corticothalamic weights substantially improved learning supported by thalamocortical weights in both tasks (**Figure 2C**). We quantified the task performance error as 1–*R*^2^ where *R*^2^ denotes the fraction of variance in the target that is predicted by the readout at the end of thalamocortical learning. The error dropped several fold across both tasks when thalamocortical learning was performed using optimized corticothalamic weights as opposed to random corticothalamic weights (**Figure 2D**; Median factor of reduction in error — Motor control: 5.8, Working memory: 5.1). We found qualitatively similar results when learning in the presence of random error-feedback weights, ***B*** (Methods; Figure S2A), showing that the improvement in learning performance cannot be attributed to feedback alignment used in the meta-learning procedure.

We next sought to understand what structure in the optimized corticothalamic weights led to improved performance. We calculated the alignment *β* between the optimized corticothalamic weights and both the direction with the highest variance (principal component direction) and the direction that drives output (readout direction) (Methods; **Figure 2E**). We express the alignment in a normalized scale, (*β* − *β*_0_)/(1 − *β*
_0_) where *β*_0_corresponds to the average alignment between meta-learned corticothalamic weights and a random direction in the cortical activity space. Whereas alignment with the readout direction was significantly greater than the alignment with the leading principal component in the motor control task, this pattern was inverted for the working memory task where in fact the corticothalamic weights were more strongly aligned with the direction of the leading principal component (**Figure 2E**). Pre-aligning corticothalamic weights to the readout direction at the beginning of the meta-learning procedure did not alter these results (Figure S2B). Together, these results demonstrate a nontrivial interplay between corticothalamic structure and task demands. Specifically, corticothalamic projections promote learning of autonomous control largely by communicating the cortical output to the thalamus, while learning of working memory benefits from communicating the principal components of the cortical activity to the thalamus.

### 3.3 Subspace aligned corticothalamic connectivity

The above analyses suggest that alignment of corticothalamic weights with specific subspaces of cortical activity improves performance. To directly test whether these subspaces are able to support learning by themselves, we consider three idealized models with categorically different forms of corticothalamic connectivity that correspond to varying degree of structure (Methods; **Figure 2B**). First, we consider unstructured connectivity that carries a *random subspace* of the cortical activity. Then, we consider connectivity aligned with the leading *principal components* (PC) of cortical activity. Finally, we consider corticothalamic connectivity that is aligned with the *readout direction* and thus transmits a copy of the network output to the thalamus.

We first trained the above models separately on different tasks by assuming maximum corticothalamic compression (*M* = 1). In both autonomous motor control and working memory tasks, we found that learning thalamocortical synapses from a single thalamic neuron is sufficient to perform the task provided the corticothalamic projection onto that neuron is chosen appropriately. For autonomous control, aligning corticothalamic connectivity with the readout direction yielded a substantial improvement over other strategies (Median −log(1 − *R*^2^); Random: 1.2, PC: 1.2, Readout: 2.1; **Figure 3A**). In contrast, the working memory task benefited from aligning corticothalamic connectivity to the leading principal component of cortical activity (Random: 1.9, PC: 2.5, Readout: 1.8; **Figure 3B**). These is consistent with the structure of corticothalamic weights optimized by meta-learning of each task (**Figure 3E**).

**Figure 3:**
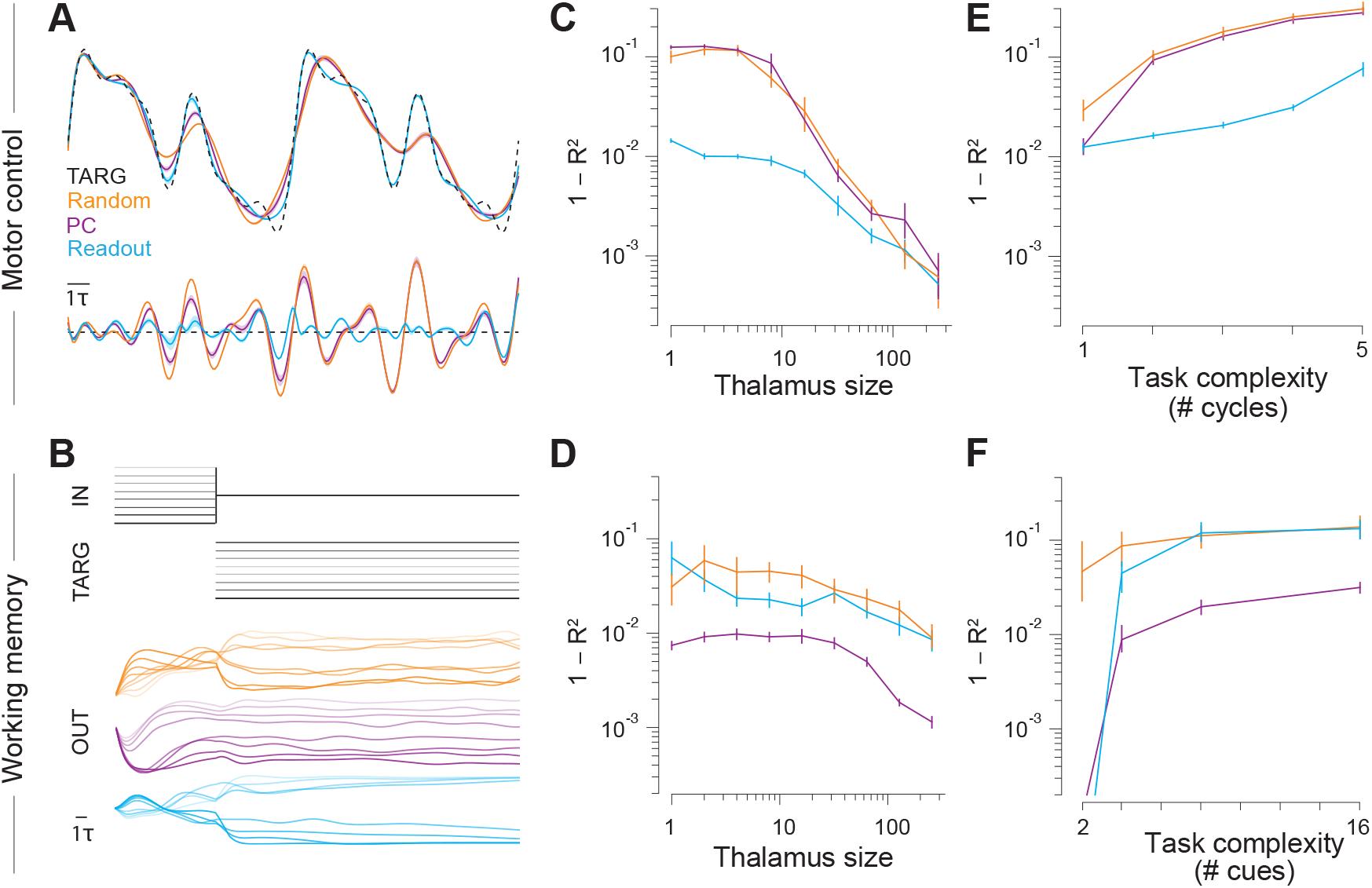
Different corticothalamic structures support different tasks. **A**. Top: Median outputs (across simulations) of networks with a single thalamic unit (*M* = 1) and different corticothalamic connectivity, trained on the autonomous control task. Bottom: The deviation of the output from the target function (bottom). Black dashed line denotes the target function. **B**. Network inputs and outputs for 8 different conditions of the working memory task. **C**. Median performance of models with different numbers of thalamic units, *M*. Lower values of 1 − *R*^2^ correspond to better performance. **D**. Similar to **C**, but for the working memory task. **E**. Median performance as a function of task complexity (Methods). **F**. Similar to **E**, but for the working memory task. All models have *N* = 256 cortical units. Error bars are standard errors.

Models with a single thalamic neuron represent an extreme scenario in which only a one-dimensional subspace of cortical activity supports learning. We therefore asked how our results change for higher-dimensional corticothalamic projections. We tested this by training the above models for a range of thalamic sizes (*M*). When aligning corticothalamic connectivity with the readout subspace, the remaining *M* − 1 thalamic neurons receive random projections. When corticothalamic connectivity is aligned with PCs, each thalamic neuron receives one of the *M* leading PCs of cortical activity. As expected, the performance of all models gradually improved with the number of thalamic neurons (**Figure 3C,D**). This makes sense because an increasingly larger subspace of cortical activity participates in learning, as we approach the limit where there would be no compression (*M* = *N*). Nonetheless, the advantage of aligning corticothalamic projections with the readout direction (for autonomous control; **Figure 3C**) or principal component directions (for working memory; **Figure 3D**) persists even when *M* is relatively large, demonstrating that structured connectivity is beneficial for realistic ratios of thalamic to cortical neurons.

To verify that the relative performance of different corticothalamic structures is governed by the type of task and not the level of complexity, we trained the models on variants of both types of tasks at different levels of complexity. We varied task complexity by either scaling the frequency of the waveform (autonomous control task), or increasing the number of unique input amplitudes to memorize (working memory task). We found that the choice of corticothalamic structure that maximized learning is robust to task complexity, depending only on the type of task being learned (**Figure 3E,F**). An exception is when the tasks are made too simple, where performance is nearly perfect regardless of whether the corticothalamic structure is aligned with the principal component direction or the readout direction. Notably, even for the simplest task variants, models with random corticothalamic connectivity learn poorly.

To understand why the relationship between corticothalamic structure and learning performance is task dependent, we examined the principal components of the cortical dynamics in both tasks. In the autonomous motor control task, the temporal fluctuations of the leading principal components tend to be very slow (Figure S3G). Communicating signals in this subspace to the thalamus cannot support learning because these signals are not adequate to generate the high frequency components in the target function. In contrast, the principal components of the cortical activity in the working memory task are dominated by the transient external inputs, and therefore encode the identity of the input (Figure S3H). Projecting these signals to the thalamus allows thalamocortical synapses to facilitate learning by contributing to building persistent activity that encodes stimulus identity during the subsequent delay period.

Taken together, these results highlight the need for communicating signals from specific subspaces via corticothalamic projections and suggest that the subspace of cortical activity to which thalamic neurons should be tuned is task-dependent. The benefits of optimized corticothalamic connectivity are not observed unless thalamocortical synapses are learned (Figure S3A,B). This suggests that performance cannot be improved if thalamus simply relays the activity in this subspace. Instead, structured corticothalamic connectivity improves task performance by enabling thalamocortical learning. Furthermore, we found that the readout signal need not be concentrated in a single thalamic neuron in order to support learning motor control (Figure S3C). Likewise, for the working memory task, different principal components need not be segregated in different thalamic neurons (Figure S3D). Therefore, thalamocortical learning can benefit from specific corticothalamic connectivity even if signals conveyed to individual thalamic neurons are not perfectly aligned with the readout or principal components. Finally, we found that structure in the corticothalamic connectivity is useful even if this structure is established gradually in conjunction with the learning of thalamocortical synapses (Figure S3E,F). Thus learning is facilitated regardless of whether the corticothalamic connectivity is developmentally hardwired (as for efferent copies of signals to the motor periphery) or learned via activity-dependent mechanisms (as for projections of the principal components of cortical activity).

### 3.4 Composite task: Delayed reaching

We have seen that different patterns of corticothalamic compression benefit autonomous control and working memory. However, naturalistic tasks often require both computations. To test whether our results generalize to such settings, we consider the problem of goal-directed reaching inspired by experiments in primates (Churchland et al., 2012). In this task, the model is required to execute reaching movements to a transiently cued location in a two-dimensional space represented on a screen. However, the movement can be initiated only upon the arrival of a go cue which follows the location cue after a random delay. Although the learning objective is solely a function of action and does not explicitly promote working memory, success in this task depends both on the ability to remember the target location during the delay (working memory) and the ability to execute movements in the absence of visual feedback (autonomous control). Therefore, we hypothesized that performance would be improved when different cortical subspaces are communicated to the thalamus during the delay and movement periods. Specifically, we hypothesized that learning benefits from communicating the principal component directions during the delay period versus the readout direction during the movement period.

To test this, we considered a thalamocortical model in which a cortical network is reciprocally connected to two distinct thalamic nuclei that are active either during preparation or execution (Methods; **Figure 4A**). This architecture is motivated by recent experiments which show that distinct populations of thalamic neurons are active before and during movement (Gaidica, Hurst, Cyr, & Leventhal, 2018). Due to the two-dimensional nature of this task, we considered a minimal model in which thalamic nuclei have two neurons each (*M* =2). The cortex receives transient input pulses whose amplitudes encode one of eight possible target locations and a binary input which serves as the go signal. The goal is to generate a pair of temporal waveforms that correspond to torques applied to the links of a two-joint arm, such that the endpoint of the arm reaches the target location (Methods). We systematically varied the structure of corticothalamic connectivity onto both thalamic nuclei to select between a random subspace, leading principal components, or read-out directions. Thalamocortical synapses were trained with RFLO. Consistent with our hypothesis, maximal performance was obtained in a model that conveyed principal components and readout signals to the thalamic nucleus that engaged in learning during preparation and execution, respectively (**Figure 4B**). This model exhibited excellent reaching performance to all targets (**Figure 4C**). The results are robust both to the number of target locations and the distribution of delays (Figure S4). These results suggest that thalamocortical learning allows the cortex to perform different movements over a range of realistic delays, provided the corticothalamic structure is chosen appropriately.

**Figure 4:**
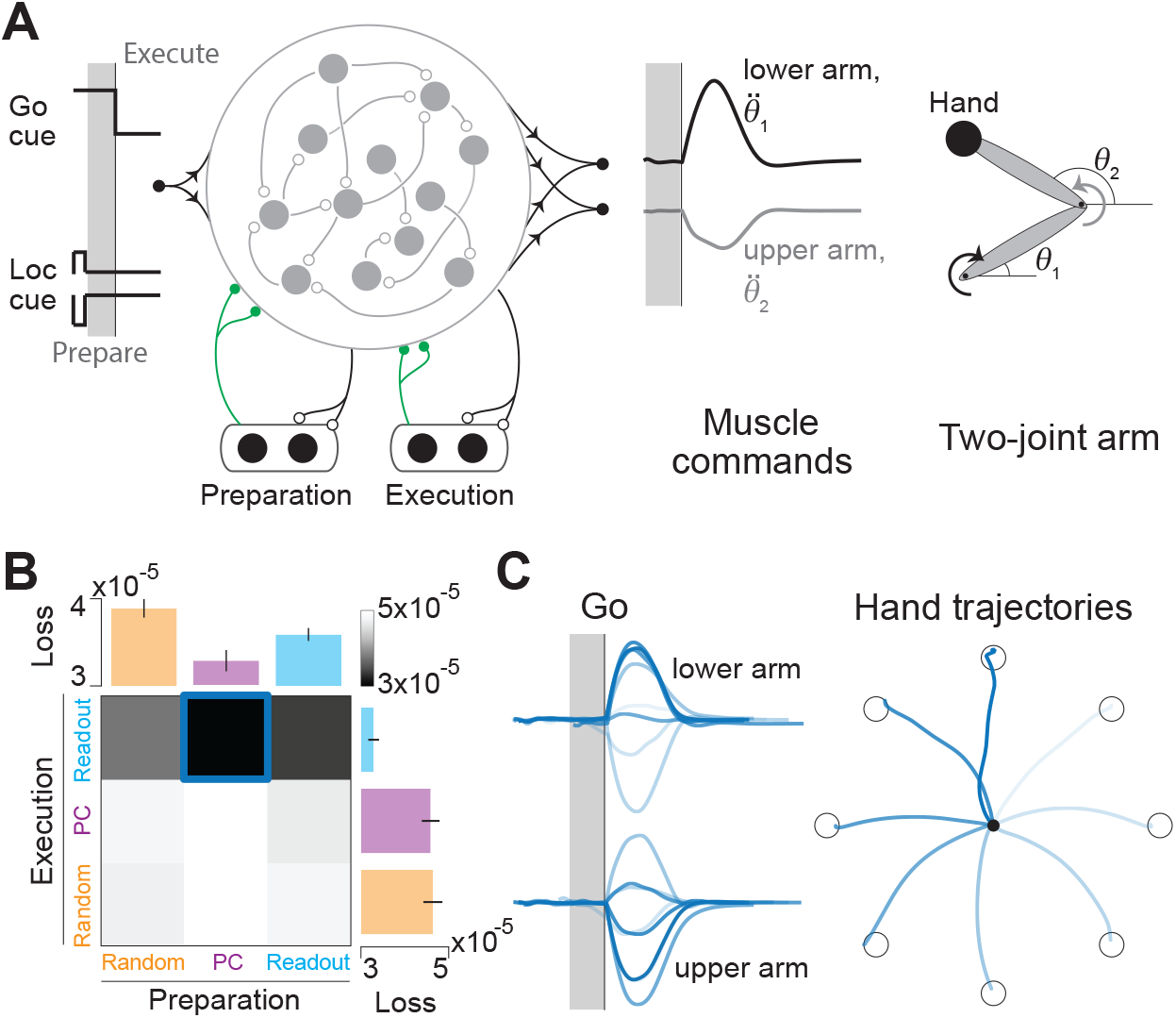
Thalamocortical model of goal-directed reaching. **A**. The model comprises two thalamic modules, one of which is active only during movement preparation and the other only during movement execution (dark gray period). The 2D output of the network controls the angular accelerations of the links of a two-joint arm. **B**. Performance of models with different types of corticothalamic connectivity onto preparatory and execution thalamic modules. Median performance of the best models from each row and column are shown in the bar plots. **C**. Left: Outputs of the best model for 8 different reach conditions, shown in different shades of gray. Right: Hand trajectories (center-out) corresponding to the outputs shown on the left. Open circles show target locations. Error bars denote standard errors.

### 3.5 Data are consistent with model predictions

Two recent studies in rodents used a combination of neurophysiology and optogenetics to demonstrate that dexterous movement generation (Sauerbrei et al., 2020) and working memory (Guo et al., 2017) both depend on interactions between thalamus and cortex. We wanted to know whether this dependence is consistent with structured corticothalamic interactions that are optimized for learning. Since our findings indicate that thalamocortical learning of movement and memory are optimized by distinct patterns of corticothalamic interactions, we reanalyzed data from both experiments to directly test whether thalamic activity during those tasks depends on the components of cortical activity predicted by our model.

In the first task (Sauerbrei et al., 2020), neural recordings were performed in the motor cortex and motor thalamus while mice performed a reach-to-grasp movement to grab a food pellet (**Figure 5A** left, Figure S5A). In the second (Guo et al., 2017), recordings were performed in the frontal cortex specifically anterior lateral motor cortex (ALM), and thalamus – specifically ventral medial (VM) and ventral anterior–lateral (VAL) nuclei, while mice performed a delayed discrimination task to report the location of an object following a delay of ∼1.3 seconds after the object was removed (**Figure 5B** – left, Figure S5B). We used a linear regression model to decode behavior (hand acceleration or choice depending on the task) and the activity of individual thalamic neurons, from the cortical population activity (**Figures 5A-B – right**, Methods). This technique identifies which modes of cortical activity propagate to behavior (i.e. estimate of readout weights, **W**°) and to the thalamus (estimate of corticothalamic weights, **W**^TC^). We restricted our analyses to the movement period and delay period for the motor control and working memory task, respectively.

**Figure 5:**
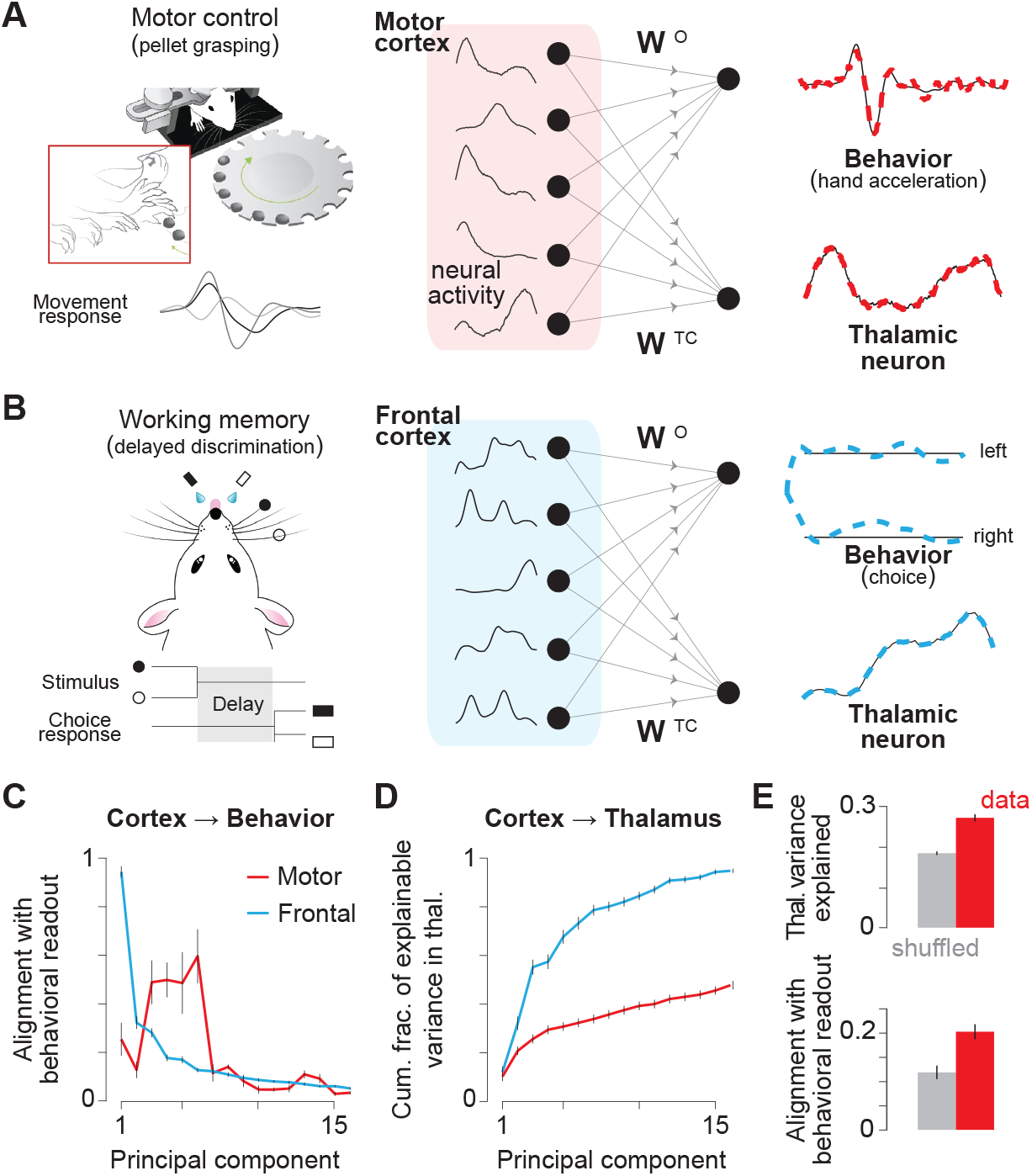
Corticothalamic interactions in mice are consistent with the model. **A**. Left: Motor control task – mice reached for a pellet of food following an acoustic cue, during recordings from motor cortex and motor thalamus. Right: Regression of cortical activity against behavior (hand acceleration) and thalamic neuron activity, respectively. Traces show the activity of a subset of cortical neurons (middle panel), and decoded estimates of behavior and example thalamic neuron activity (right panel; data in black and decoded estimate in color). Data were aligned to movement onset prior to regression. **B**. Left: Working memory task (modified from (Bjerre & Palmer, 2020)) mice reported the location of a pole by directional licking after a delay period, during recordings from frontal cortex and thalamus. Right: Regression of cortical activity against behavior (choice), and thalamic neuron activity. Data are aligned to the onset of delay period. **C**. Mean alignment between the direction of readout weights and different cortical PCs, in both tasks. Error bars denote ±1 SEM. (Motor control: n=3 sessions, Working memory: n=5). **D**. Cumulative fraction of explainable variance in thalamic activity as a function of the number of cortical PCs. Fractions were normalized to a scale between 0 and 1 for each thalamic neuron before averaging. **E**. Top: Fraction of variance in an average thalamic neuron explained by cortical PCs during the motor task. Bottom: Average alignment between readout weights and corticothalamic weights across thalamic neuron. Gray bars denote chance levels obtained by shuffling trial indices. Error bars in D and E denote standard errors in mean (Motor control: n=101, Working memory: n=72).

We found that behavior was well explained by cortical activity in both tasks (*R*^2^ — Motor control: 0.93, Working memory: 0.97). To determine specifically which directions of cortical activity correlate with these behaviors, we examined the alignment of the readout weights with the principal components of cortical activity. In mice performing the working memory task, leading principal components (PCs) of the frontal cortex activity had a large influence on the animal’s behavioral choice (**Figure 5C – blue**). In contrast, hand movements during the motor task were primarily influenced by PCs of lower variance in the motor cortex (**Figure 5C – red**, Figure S5C). This difference is consistent with our model results and suggests that the principal components of the cortex that drive behavior are different for the two tasks.

Due to the greater contribution of lower variance PCs to the behavioral readout in the motor control task, we can test the key model prediction pertaining to the structure of corticothalamic connectivity – namely whether the structure is optimized for learning each task. Specifically, we predict that corticothalamic weights should be aligned with readout weights in the motor control task. This means that we would need lower variance PCs of cortical activity to capture the response of neurons in the thalamus. In contrast, in the working memory task, we predict that corticothalamic weights should extract the leading PCs of cortical activity. If this is the case, we should be able to capture thalamic responses using only the high variance PCs. We first quantified the fraction of variance explained (*R*^2^) in each thalamic neuron when using the activity of all neurons recorded in the cortex (Methods). We found that we could capture a substantial fraction of thalamic variance in both tasks, although the fraction was higher in the working memory task (Motor control: 0.46 ± 0.01, Working memory: 0.70 ± 0.01; Figure S5D). We then repeated this analysis to compute the fraction of variance explained in each thalamic neuron when using only the *k* leading PCs of cortical activity (*R*^2^(*k*)). To compare corticothalamic interactions across the two tasks, we computed the fraction of *explainable* variance as a function of the number of PCs, *R*^2^(*k*)*/R*^2^. Consistent with the model prediction, we found that more principal components are needed to explain thalamic activity during the motor control task than in working memory task (**Figure 5D** – red vs blue, Figure S5E). The fraction of explainable variance in thalamus captured by the top 5 cortical PCs was significantly greater during the working memory task than motor control (Motor control: 0.29 ± 0.02, Working memory: 0.63 ± 0.04).

Further analysis suggested that corticothalamic interactions identified above are not solely due to common input to cortex and thalamus. In the working memory task, inhibiting cortex dramatically reduces variability in thalamus indicating that the interactions are causal (Guo et al., 2017). Similar inactivation experiments were not performed in the motor control task, but we took advantage of the fact that neural recordings were simultaneous to test whether trial-by-trial fluctuations in motor cortex propagate to the motor thalamus. We found that thalamic activity on any given trial was substantially better predicted by cortical activity in the same trial (**Figure 5E** – top) suggesting that the corticothalamic interactions do not merely reflect common inputs to motor cortex and thalamus that are similar across trials, such as context signals. Furthermore, we found that corticothalamic weights that capture trial-by-trial activity in thalamic neurons were better aligned with the readout weights than those that capture only trial-averaged thalamic activity (**Figure 5E** – bottom). Therefore, fluctuations in corticothalamic signals across trials reflect fluctuations in the output of the motor cortex, suggesting that motor thalamus receives an efference copy of the motor command.

## 4 Discussion

We investigated how the connectivity of cortico-thalamo-cortical loops contributes to learning of motor and cognitive functions. Current theories that propose a computational role for these loops (Kao et al., 2021; Logiaco et al., 2021) do not address how the connectivity weights could be learned. We found that local feedback-driven learning algorithms, which have been proposed as biologically plausible solutions for credit assignment in recurrent networks (Miconi, 2017; Gilra & Gerstner, 2017; Alemi et al., 2018; Murray, 2019) are effective for updating thalamocortical synapses but ineffective for corticothalamic synapses, challenging theories proposing a computational role for CTC loops. We resolved this challenge by developing a model in which structured corticothalamic connectivity supports thalamocortical learning. Such structure dramatically enhances learning of thalamocortical synapses, and the structure that optimizes performance depends on the task. Specifically, autonomous motor control requires a specialized corticothalamic pathway that communicates the efferent copy of the motor command, whereas connectivity structure that projects the principal components of cortical activity facilitates working memory tasks. We analyzed neural recordings from cortex and thalamus of mice during both types of tasks and found that the influence of cortex on thalamus is qualitatively consistent with these predictions.

### 4.1 Constraints on anatomy

We used meta-learning or “learning to learn” (Wang, 2021; Hospedales et al., 2022) to optimize corticothalamic weights while a local plasticity rule was applied to thalamocortical synapses. Similar approaches have recently been applied to other neural systems (Wang et al., 2018; Jiang & Litwin-Kumar, 2021; Tyulmankov, Yang, & Abbott, 2022). Since this approach involves a slow optimization process based on gradient descent, in the context of our model the structure of the corticothalamic pathway identified in this manner is best viewed as a product of evolutionary and developmental processes. This view is consistent with a lack of clear evidence for plasticity in the corticothalamic pathway during learning, and supported by our finding that error-driven, biologically plausible learning algorithms are ineffective in adjusting corticothalamic synapses. While synapses in the cortex might undergo more sophisticated forms of plasticity by leveraging the complex dendritic machinery involving distinct apical and basal compartments (Urbanczik & Senn, 2014; Guerguiev et al., 2017), such structures are absent in the thalamus and therefore cannot be used to perform credit assignment in corticothalamic synapses. It is possible that there are feedback pathways communicating error signals to the thalamus that are themselves optimized by evolution and confer corticothalamic synapses with the potential for error-driven plasticity. Modeling studies have shown that such an approach is effective in the context of feedforward networks (Lee, Zhang, Biard, & Bengio, 2014; Akrout, Wilson, Humphreys, Lillicrap, & Tweed, 2019; Lindsey & Litwin-Kumar, 2020), and would make for an interesting alternative hypothesis of learning in thalamocortical loops. However, the predictions of such a model would be difficult to test because to our knowledge, pathways conveying error signals to the thalamus, if any, are yet to be mapped out. Instead, we optimized corticothalamic synapses to make predictions that can be corroborated with available anatomical and physiological data.

Anatomical tracing studies show that there is, indeed, an evolutionarily conserved pathway from layer V pyramidal neurons in the cortex to higher-order thalamus through axonal branching of corticofugal projections (Bourassa, Pinault, & Deschênes, 1995; Bourassa & Deschênes, 1995; Kakei, Na, & Shinoda, 2001; Kita & Kita, 2012; Sherman, 2016). Since a major target of corticofugal projections is lower motor centers such as the spinal cord, this pathway is ideally suited to convey an efference copy of the cortical motor output (Guillery & Sherman, 2011; Sherman & Usrey, 2021). This is precisely the signal that would optimize thalamocortical learning of motor control according to our model. Our analysis of neural data confirmed that the alignment of the activity of thalamic neurons with the cortical motor output was significantly greater than chance during pellet grasping. While the degree of alignment was not perfect, we showed that learning with partial alignment is sufficient to facilitate learning (**Figure S3C**). It should be noted that we used acceleration of the limb as a proxy for motor output, which is not perfectly correlated with EMG signals (Keil, Elbert, & Taub, 1999; Fougner, Scheme, Chan, Englehart, & Stavdahl, 2011; Chen et al., 2011). Furthermore, one recent study showed that corticofugal projections that branch into the thalamus primarily target midbrain and not spinal cord (Economo et al., 2018), suggesting that thalamocortical loops might not participate in learning all types of movement. Determining the conditions under which neural activity in cortico-recipient motor thalamus resembles an efference copy measured using EMG signals will help elucidate the function of thalamus in motor learning.

Our model also suggests that a motor efference copy is not required for all types of tasks. For learning to maintain stimulus identity in working memory, communicating the principal components of cortical activity to the thalamus is sufficient. This could be accomplished through unsupervised plasticity mechanisms that tune corticothalamic weights. Unlike error-based learning, unsupervised mechanisms such as Hebbian learning do not require credit assignment and can successfully operate on corticothalamic synapses (Magee & Grienberger, 2020). This possibility is consistent with our analysis of neural data during the delayed discrimination task which suggests that the corticothalamic pathway from ALM to VM/VAL complex communicates the principal components of activity in ALM. While there is evidence for both Hebbian and homeostatic forms of plasticity in sensory thalamus (Krupa, Ghazanfar, & Nicolelis, 1999; Butts, Kanold, & Shatz, 2007; Krahe & Guido, 2011; Taylor et al., 2021), evidence for plasticity in higher-order thalamus is limited (Ding, Li, Clark, Diaz, & Rafols, 2003). Our results identify the pathway from ALM to thalamus (Guo et al., 2018) as a leading candidate to investigate whether corticothalamic synapses undergo Hebbian plasticity, but we note that such mechanisms may be at play even in thalamic nuclei receiving motor efference copies.

### 4.2 Assumptions and extensions

In our models, thalamocortical synapses are updated according to a local plasticity rule for error-based learning (Murray, 2019). There is also evidence for a role of dopamine in mediating cortical plasticity (Hosp, Pekanovic, Rioult-Pedotti, & Luft, 2011; Li et al., 2017) suggesting that reward-based learning rules may also modify thalamocortical synapses. Further research is needed to verify whether our results generalize to other forms of plasticity including spike-based algorithms (Gilra & Gerstner, 2017). Furthermore, we used a coarse-grained model composed of a homogeneous neural populations in the cortex and thalamus. However, recent experiments show evidence for convergent inputs from multiple cortical areas to individual thalamic neurons (Sampathkumar, Miller-Hansen, Sherman, & Kasthuri, 2021), rich heterogeneity in wiring across cortical layers that project to/from thalamus (Munñz-Castañeda et al., 2021), and diversity in the membrane properties across thalamic neurons (Phillips et al., 2019), placing important constraints on computation in CTC loops. Furthermore, thalamus receives commands from other brain structures like the cerebellum and BG which may interact with corticothalamic inputs to influence thalamocortical plasticity (Tanaka et al., 2018). Corticothalamic weights in our model contain both positive and negative values. Although we do not explicitly model excitatory and inhibitory populations, negative weights can be implemented by feedforward inhibition from cortex to thalamus via the thalamic reticular nucleus (TRN). TRN also mediates local inhibition in the thalamus, which we did not include in our models as it is beyond the scope of this study.

### 4.3 Relationship to other studies

Our model expands a body of work that conceptualizes connectivity in recurrent networks as a sum of a random matrix and a tunable low-rank perturbation (Mastrogiuseppe & Ostojic, 2018; Schuessler, Dubreuil, Mastrogiuseppe, Ostojic, & Barak, 2020), and is closely related to recent studies that suggest a role for thalamocortical loops in motor control. Specifically, it has been shown that CTC loops can support motor preparation by tuning the thalamocortical weights to minimize prospective errors (Kao et al., 2021) and facilitate fluid transition between learned movement motifs in addition to supporting noise-robust movement execution (Logiaco et al., 2021). In these studies, synaptic weights were set via numerical optimization without specifying how this optimal solution could be learned using biological mechanisms. By postulating that CTC loops are optimized for the task of learning to minimize errors, we show how these objectives may be achieved in a biologically plausible way. At the same time, we accommodate experimental findings which demonstrate that behavioral improvement during training is accompanied by thalamocortical plasticity (Biane et al., 2016; Audette et al., 2019; Hasegawa et al., 2020; Sohn et al., 2022).

Our results also suggest that changes in cortical representations observed during learning (Huber et al., 2012; Peters, Chen, & Komiyama, 2014; Masamizu et al., 2014) may not be entirely due to plasticity of corticocortical (CC) synapses. There is evidence for corticocortical plasticity which contributes to activity remapping observed during training (Biane, Scanziani, Tuszynski, & Conner, 2015; Biane, Takashima, Scanziani, Conner, & Tuszynski, 2019; Hasegawa et al., 2020). However, we found that CTC loops can contribute substantially even in the presence of corticocortical plasticity. Determining the differential contribution of thalamocortical and corticocortical plasticity to learning is an important direction for future studies. One possibility is that different CTC loops are engaged in different contexts such that thalamocortical plasticity contributes to learning context-specific behavioral components whereas corticocortical plasticity helps consolidate components that are shared across contexts. Such coordination between cortex and thalamus is a potential solution to the problem of continual learning (Wang & Halassa, 2022). A related proposal is that corticothalamic synapses facilitate transfer of control over flexible sensorimotor associations from basal ganglia (BG) to CTC loops (Wang & Halassa, 2022). Although we do not explicitly model BG, we demonstrated that our model can take advantage of thalamic control by BG to learn composite reaching tasks by switching between thalamic units with appropriate corticothalamic structures during different phases of the task (**Figure 4C**).

### 4.4 Conclusion

Task-optimized recurrent neural networks have become a leading tool to investigate neural mechanisms and representations underlying cognition and motor control. However, the standard approach of generic network architectures in which synaptic weights are trained end-to-end using, e.g., the backpropagation algorithm, is not a biologically plausible model of the learning process itself (Mante, Sussillo, Shenoy, & Newsome, 2013; Sussillo, Churchland, Kaufman, & Shenoy, 2015; Barak, 2017). Therefore, such approaches do not reveal how the brain makes use of specific anatomical circuit motifs and local plasticity rules to facilitate learning. We fill this gap in the context of cortico-thalamo-cortical loops by demonstrating that task-specific, structured corticothalamic connectivity is necessary to optimize learning when biologically plausible plasticity rules are employed, thereby establishing a link between anatomy and computation. Our findings show that understanding the structure of corticothalamic connectivity may be key to determining the computational role of higher-order thalamus in orchestrating behavior.

## 5 Methods

### Task description

We trained networks separately to solve three different types of tasks based on experiments that study computations underlying different functions – motor control, working memory, and goal-directed reaching. The motor control task tests the ability of networks to generate complex temporal patterns (like electromyograms) autonomously in the absence of external inputs. The working memory task tests how well networks maintain the identity of transient inputs, shortly after they have been removed. The reaching task tests a combination of the above two skills: the ability to remember a transiently cued target location, and then generate an appropriate movement pattern to reach that location upon the arrival of a go cue.

#### Motor control

The input **x**(*t*) is set to zero. The target function is a sum of sinusoidal functions, 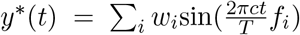, where **f** = [1, 2, 4, 6] denotes the frequencies of the sinusoids, **w** = [1, 0.75, 0.5, 0.25] denotes their relative strengths, and *T* is the total duration. In the basic version of the task, we set *c* = 2 and *T* = 20*τ* where *τ* is the time-constant of the cortical neurons. Task complexity is controlled by varying *c* from 1 to 5 which has the effect of scaling all frequency components of the target function by the same factor.

#### Working memory

The input **x**(*t*) is a pulse with one of *P* possible amplitudes *Ap*spaced evenly between +1 and −1 and a duration of 10*τ*. The goal is to reproduce the input amplitude during the subsequent delay period, so the target is a constant function *y*^*^(*t*) = *Ap*for a period of 30*τ* after the end of the input pulse. The basic version of the task corresponds to *P* = 8. Task complexity is controlled by varying *P* from 1 to 16 on a logarithmic scale.

#### Reaching

The input **x**(*t*) ∈ ℝ^2^ comprises the target location encoded in the amplitude of a 2D pulse of duration 4*τ*, and the go signal encoded in a step function 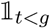 where *g* ∈ (5*τ*, 20*τ*) denotes the variable timing of the go signal. The goal is to generate a 2D output pattern that is zero for *t < g*, and subsequently deliver appropriate torques to control a two-link arm, such that the endpoint of the arm reaches the target location within a period of 10*τ*. The moment of inertia of the arms was taken to be unity such that the amount of torque applied is identical to the angular acceleration of the arms. The precise temporal pattern of angular acceleration, **y**^*^(*t*) needed to perform successful reaching was obtained by training a recurrent neural network via backpropagation through time. The function **y**^*^(*t*) was then used as the target function for training biologically plausible models.

### Thalamocortical model

The thalamocortical model has a network architecture in which *N* interconnected cortical neurons is reciprocally connected with *M* uncoupled thalamic neurons where *M* ≪ *N*. In this model, the cortical population activity **h** depends on thalamic activity **r** and external input **x** according to Equation 1, 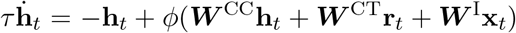, where the nonlinearity *ϕ*(·) is the tanh function. The thalamic activity depends only on cortical activity as there is no recurrence within the thalamus, **r**_*t*_= *ϕ*(***W*** ^TC^**h**_*t*_), and the network output is assumed to be a linear readout of the cortical activity: **y**_*t*_= ***W*** °**h**_*t*_. Simulations were performed using a timestep, dt = 10ms, and the neuronal time constant was set to *τ* = 100ms (10 timesteps). The dimensionality of the network input, *S*, and the network output, *R*, are task specific. *S* is 0,1, and 3 respectively for motor control, working memory, and reaching. *R* = 1 for motor control and working memory tasks and *R* = 2 for reaching. The number of cortical units, *N*, is fixed at 256 and the number of thalamic units, *M*, is varied systematically from 1 to 256 on a logarithmic scale. Elements of the input weight matrix ***W*** ^I^and the recurrent weight matrix ***W*** ^CC^are Gaussian, sampled from 𝒩(0, 1*/S*) and 𝒩(0, *g*^2^*/N*), respectively. The strength of recurrent connectivity was chosen such that the network operated in the chaotic regime, *g* = 1.5. Output weights ***W*** °are sampled from a uniform distribution, 𝒰(0, 2*/N*). Thalamocortical weights ***W*** ^CT^are sampled from 𝒩 (0, *g*^2^*/M*).

Connection sparsity in the cortex is controlled by varying the fraction of cortical neurons that each cortical neuron makes a synaptic contact with (*f*). The fraction is varied from *f* = 1*/N* (very sparse) to *f* = 1 (fully connected) on a log scale. To ensure that the variance of total input current into single neurons remains independent of connection sparsity, we sampled the elements of ***W*** ^CC^from 𝒩(0, *g*^2^/(*f N*)). The dynamical regime of the cortex is controlled by varying the strength of recurrence within the cortex, *g* = [0.5, 0.75, 1.0, 1.25, 1.5]. For the simulations in which we vary the output dimensionality, we use variants of the motor control and working memory tasks with *R* = [1, 2, 3, 4] outputs.

### Canonical models of corticothalamic connectivity

We consider three types of thalamocortical models that differ in the structure of corticothalamic connectivity.

#### Random

In this version of the model, elements of ***W*** ^TC^are sampled randomly from 𝒩(0, 1*/N*). Therefore, the thalamic activity in this model corresponds to a random low-dimensional projection of the cortical activity. These weights are held fixed throughout learning.

#### Principal component

In this strategy, elements of ***W*** ^TC^are proportional to the *M* leading eigenvectors *U* of the covariance in cortical population activity, where the constant of proportionality is 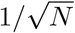. The resulting thalamic activity corresponds to the *M* leading principal components of the cortical activity. Since the structure of cortical dynamics changes during thalamocortical learning, corticothalamic weights are dynamically updated in this version at the end of each learning trial: 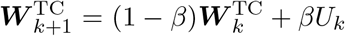 where *U* _*k*_ denotes the leading eigenvectors of the cortical covariance at the end of *k* trials, and *β* = 0.01 is the corticothalamic learning rate. Biologically, the dynamic updating of corticothalamic weights in this manner corresponds to a Hebbian learning strategy operating at the corticothalamic synapses. We also trained a variant of this strategy where individual principal components are distributed across all *M* thalamic neurons instead of being segregated in individual neurons. This variant was constructed by multiplying the corticothalamic weights that yield segregated PCs, by a random orthonormal matrix.

#### Readout

This version of the model is characterized by a perfect alignment between readout weights ***W*** °and corticothalamic weights onto a small subset of the thalamic units. This subset contains a maximum of *R* units whose activity mirrors the network output, where *R* denotes the number of output units. Corticothalamic projections onto the remaining *M* − *R* thalamic units are sampled randomly from 𝒩(0, 1*/N*). Similar to the principal component strategy, the corticothalamic weights onto the subset of *R* thalamic neurons are updated dynamically: 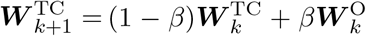 where 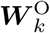 denotes the readout weights at the end of *k* trials and *β* = 1.

Biologically, this strategy is analogous to a pathway that carries signals from cortex to thalamus via axon collaterals. For models with *M > R*, we also trained a variant of this strategy where the *R*-dimensional readout signal is distributed across all *M* thalamic neurons instead of being concentrated in *R* neurons. This variant was constructed by multiplying the corticothalamic weights by a random orthonormal matrix.

### Learning algorithm

Thalamocortical weights are trained using a biologically plausible algorithm called Random Feedback Local Online, RFLO (Murray, 2019) (Figures 2-4). With the exception of Figure 1C (see below), we use meta-learning to optimize corticothalamic weights while learning thalamocortical weights concurrently via RFLO (Wang, 2021; Hospedales et al., 2022)(Figure 2).

#### RFLO

The goal of learning is to minimize the discrepancy between the network output and a desired target function. Therefore, the loss function is taken to be the squared-error across all output units, integrated over time where the error in dimension *r* of the output is the difference between the target and the actual output of the network in that dimension, 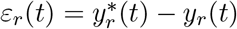.

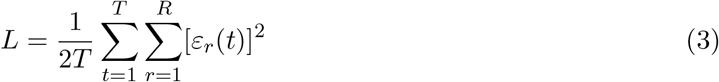

The weight updates in RFLO are derived by making two approximations to standard gradient-based algorithms that minimize the loss with respect to the weights. The first approximation consists of dropping nonlocal terms from the gradient, so that computing the update to a given synaptic weight requires only pre- and postsynaptic activities, rather than information about the entire state of the cortex including all of its synaptic weights. Second, we project the error back into the cortical network for learning using random feedback weights ***B*** sampled from 𝒰(0, 2*/N*), rather than feedback weights that are precisely tuned to match the readout weights. This relaxation is made possible by learning readout weights in conjunction with thalamocortical weights. Third, the weight updates are performed in real-time instead of accumulating gradients and updating the weights at the end of each trial. The resulting update rules for thalamocortical weights ***W*** ^CT^and readout weights ***W*** °are given by:

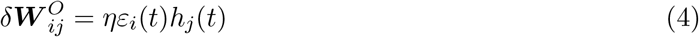

where *h* _*j*_is the activity of cortical unit *i*, 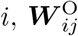 denotes the readout weight from cortical unit *j* to output unit *i*, and *η* is the learning rate.

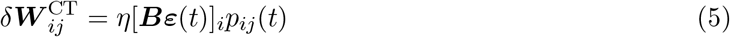

where 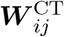 denotes the thalamocortical weight from thalamic unit *j* onto cortical unit *i*; *p*_*ij*_ denotes the eligibility trace that accumulates the correlation between the activity of the (presynaptic) thalamic unit *j* and the (postsynaptic) cortical unit 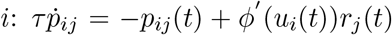 where *r* _*j*_ is the activity of thalamic unit *j*, 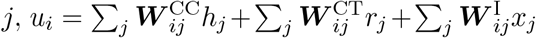 is the total input current to cortical unit *i*, and *α* is the leak rate of the cortical units. For the simulation in Figure 1C, the update rule for the update rule for corticocortical weights ***W*** ^TC^is given by:

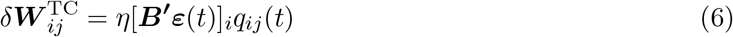

where 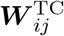 denotes the corticothalamic weight from cortical unit *j* onto thalamic unit *i*; *q* _*ij*_denotes the eligibility trace defined as the product of the activity of the (presynaptic) cortical unit *j* and the (postsynaptic) thalamic unit 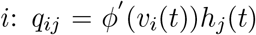 where *h* _*j*_ is the activity of cortical unit *j*, 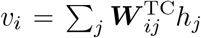 is the total input current to thalamic unit *i*. ***B***^***′***^denotes the random feedback weights through which error signals arrive at the thalamus. Elements of ***B***^***′***^are sampled from 𝒰(0, 2*/M*). Prior to learning, thalamocortical weights and readout weights are both initialized randomly by sampling from 𝒩(0, *g*^2^*/M*) and *U* (0, 2*/N*) respectively. The learning rate was set to *η* = 0.1 for readout and thalamocortical weight updates, and the training was performed for *K* = 10, 000 trials unless specified otherwise. The learning rate for corticothalamic weights for the simulation in Figure 1C was determined by hyperparameter optimization (*η* = 0.001).

#### Meta-learning

We use meta-learning to determine the optimal structure of corticothalamic projection that supports biologically plausible of thalamocortical weights. Corticothalamic weights *W* ^TC^are initialized randomly at the beginning of the meta-learning procedure by sampling from 𝒩(0, 1*/N*), and weights are updated using backpropagation through time at the end of each epoch containing *K* = 200 learning trials. The objective is to minimize the average loss 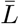 across trials of the epoch, 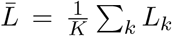 where the loss in each trial *L*_*k*_ is given by Equation 3. Thalamocortical weights *W* ^CT^are reinitialized randomly at the beginning of each epoch by sampling from 𝒩(0, *g*^2^*/M*), and weights are updated using RFLO according to Equation 5. We considered a model with *N* = 256 cortical neurons and *M* = 32 thalamic neurons. Meta-learning was used to update corticothalamic weights onto only one of the thalamic units. Optimization is performed for 10,000 epochs using Adam optimizer with a learning rate of 0.0001. By restricting meta-learning to a single thalamic unit, we can readily evaluate the relative optimality of the different strategies used in Figure 2 by measuring the alignment of the meta-optimized corticothalamic weights with the readout direction and the principal component direction.

### Statistical analyses

Each model tested in this study was simulated 40 times with different parameter initializations, and those initializations were identical across models sharing similar architecture. Unless specified otherwise, we used median as the summary statistic and error bars denote standard errors estimated by bootstrapping.

#### Task performance

We quantify task performance as 1 − *R*^2^ where 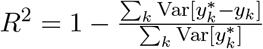 and Var[·] denotes variance across time. For tasks with multiple conditions – Working memory and Reaching – the outputs from different conditions were concatenated before computing *R*^2^. A value of 0 corresponds to perfect performance, while 1 corresponds to chance level. Note that this measure reduces to the loss *L* defined in Equation 1 when the target function has unit temporal variance.

#### Alignment between weights

We quantified the alignment between pairs of weight vectors **m** and **n** by taking their normalized dot product 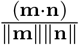 where ∥·∥ denotes the 𝓁_2_ norm. For computing feedback alignment in the cortex (Figure 1C), **m** = ***B*** and **n** = ***W*** °where ***B*** denotes the feedback weights projecting the error signal to cortical neurons. For computing feedback alignment in the thalamus (Figure 1C), **m** = ***B***^***′***^and **n** = ***W*** ^CT^*ϕ*^′^(**u**_*t*_)***W*** °where ***B***^***′***^denotes the feedback weights projecting the error signal to thalamic neurons. The alignment between optimized corticothalamic weights and readout weights is computed by taking, **m** = ***W*** ^TC^and **n** = ***W*** °. The alignment between optimized corticothalamic weights and principal component weights is computed by taking, **m** = ***W*** ^TC^and **n** = ***z***, where ***z*** denotes the leading eigenvector of the cortical covariance matrix.

#### Neural datasets

Detailed experimental methods for behavioral and neural recordings in the motor control task and working memory task are described in (Guo et al., 2015; Sauerbrei et al., 2020) and (Guo et al., 2017) respectively. In the motor control task (Sauerbrei et al., 2020), neural recordings were performed simultaneously in the motor cortex and motor thalamus while mice performed a reach-to-grasp movement to grab a food pellet (**Figure 5A** – left). The dataset includes one behavioral session each from 3 different mice. The mean number of cortical and thalamic units was 53 ± 9 and 33 ± 5 respectively. In the working memory task (Guo et al., 2017), recordings were performed in separate sessions in the frontal cortex, specifically anterior lateral motor cortex (ALM), and thalamus, specifically ventral medial (VM) and ventral anterior–lateral (VAL) nuclei, while mice performed a delayed discrimination task to detect (by whisking) and report (by licking left/right) the location of a pole (anterior/posterior) following a delay of ∼1.3 seconds after the pole was removed (**Figure 5B** – left). The dataset includes 5 behavioral sessions from one mouse. The total number of cortical and thalamic units was 151 and 72 respectively.

#### Estimation of readout weights

We estimated the readout weights for both tasks by regressing behavior against the activity of the population of cortical neurons. We split the trials into a training set (80% for estimating weights), a validation set (10% for hyperparameter optimization), and a test set (10% for computing variance explained *R*^2^).

For the motor control task, behavior was defined as the time-varying acceleration profile of the hand. To obtain acceleration profiles, hand position traces were first aligned to the onset of hand movement and averaged across trials in the training set. We then numerically computed the second derivative to obtain the average acceleration profile along the three axes (*a* _*x*_(*t*), *a*_*y*_(*t*), *a* _*z*_ (*t*)). Likewise, we aligned spike trains of motor cortex neurons to the onset of hand movement, convolved them with a gaussian function (width *σ* as hyperparameter), and computed the trial-averaged population activity, **h**(*t*) in the training set. We regressed the acceleration profiles against the population activity to estimate the readout weights 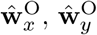 and 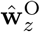 using ordinary least squares, and optimized the smoothing parameter *σ* by cross-validation. Finally, we expressed the readout weights in the basis of the principal components (PC) of the population activity. To do this, we first performed eigendecomposition of the population covariance, ⟨**hh**^T^⟩ = *U* Λ*U* ^T^, and then projected the estimated readout weights for each component of acceleration onto the top *k* = 16 eigenvectors, e.g., 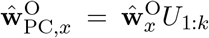. Since the weight profiles of the three components of acceleration were qualitatively similar in the PC basis, we averaged them to obtain a single readout weight profile for the motor control task 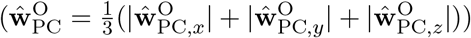.

For the working memory task, behavior was defined as the choice made on each trial. We aligned spike trains of ALM neurons to the onset of the delay period, convolved them with a gaussian function (width *σ* as hyperparameter), and averaged them separately across trials with leftward and rightward choices (**h** _*l*_(*t*) and **h** _*r*_(*t*)). We restricted out analysis to trials in which the choice was correct. We concatenated the response from the two sets of trials (**h**(*t*) = [**h** _*l*_ (*t*) **h** _*r*_(*t*)]) and used them as predictors in a linear regression model to decode choice, *c*(*t*) = [*c*_*l*_ (*t*) *c* _*r*_ (*t*)], where *c*_*l*_(*t*) = −1 and *c*_*r*_ (*t*) = +1. As in the motor control task, readout weights **Ŵ**°were estimated using ordinary least squares, and we optimized the smoothing parameter *σ* by cross-validation. Finally, we expressed the readout weights in the basis of the principal components (PC) of the population activity to obtain 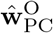.

#### Analysis of corticothalamic communication

For estimating corticothalamic weights, we used a procedure similar to the one outlined above, except now the activity of cortical neurons **h**(*t*) was used to decode the activity of individual thalamic neurons *r*(*t*), instead of decoding behavior. We used the principal components of cortical activity as predictors instead of raw firing rates. Specifically, we first performed eigendecomposition of the population covariance, ⟨**hh**^T^⟩ = *U* Λ*U* ^T^, and projected the cortical activity onto the top *k* eigenvectors, **h** _*k*_ (*t*) = **h**(*t*)*U* 1:_*k*_. We then estimated the regression weights to decode *r*(*t*) from **h** _*k*_ (*t*). By varying *k* from 1 to 16, we estimated the corticothalamic weights 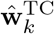 that captured the influence of the top *k* principal components of cortical activity on each thalamic neuron. The cumulative variance in the thalamic neuron activity explained by the top *k* cortical principal components was quantified using the coefficient of determination, 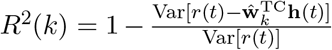. Finally, we divided the cumulative variance explained by *k* principal components by the variance explained all *K* = 16 principal components to obtain a normalized measure of variance explained, *R*^2^(*k*)*/R*^2^(*K*), for each thalamic neuron in the dataset.

