## supplemental figures for "Specific connectivity optimizes learning in thalamocortical loops"

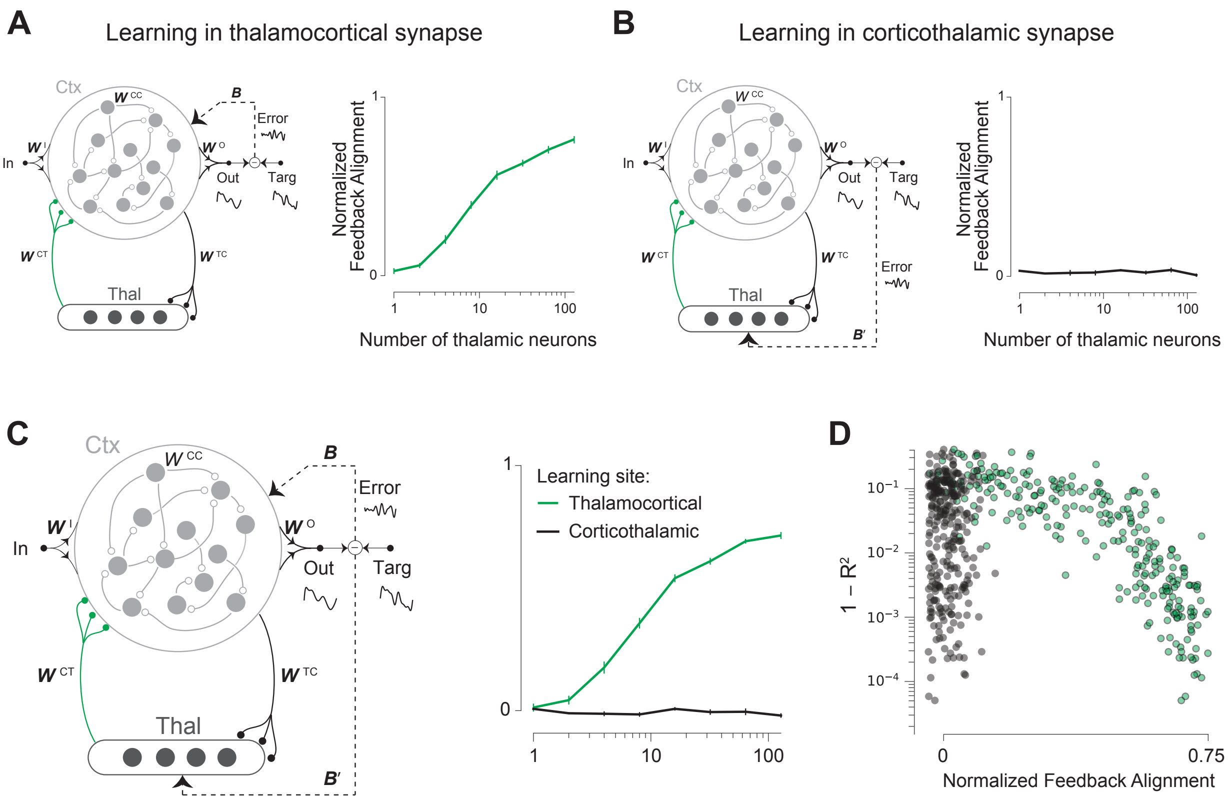


Figure S1: **Local plasticity of thalamocortical and corticothalamic synapses**. **A-B.** Left: Corticocortical weights ($\boldsymbol{W}^{\mathrm{CC}}$) are fixed. A local learning rule (RFLO, see text) is applied to update thalamocortical weights ($\boldsymbol{W}^{\mathrm{CT}}$) in **A** or corticothalamic weights ($\boldsymbol{W}^{\mathrm{TC}}$) in **B**. Error signals are projected to the cortex (via $\boldsymbol{B}$) and thalamus (via $\boldsymbol{B'}$) to facilitate learning in thalamocortical and corticothalamic synapses respectively. Right: The alignment between feedback weights ($\boldsymbol{B}$) and readout weights ($\boldsymbol{W}^{\mathrm{O}}$) in cortical neurons (**A**). Alignment between feedback weights ($\boldsymbol{B'}$) and effective readout weights ($\boldsymbol{W}^{\mathrm{CT}}\phi'(\mathbf{u}_t)\boldsymbol{W}^{\mathrm{O}}$, see Methods) in thalamic neurons (**B**). **C.** Left: RFLO is applied to simultaneously update both thalamocortical weights and corticothalamic weights. Right: The alignment between feedback weights ($\boldsymbol{B}$) and readout weights ($\boldsymbol{W}^{\mathrm{O}}$) in cortical neurons increases as learning performance improves (see Figure 1C – black) indicating successful credit assignment in thalamocortical synapses (green). In contrast, there is no alignment between feedback weights ($\boldsymbol{B'}$) and effective readout weights ($\boldsymbol{W}^{\mathrm{CT}}\phi'(\mathbf{u}_t)\boldsymbol{W}^{\mathrm{O}}$, see Methods) in thalamic neurons indicating the failure of credit assignment in corticothalamic synapses (black). Numerical estimates of alignment ($\beta$) are normalized to correct for the chance level alignment ($\beta_0$) before plotting, $(\beta - \beta_0)/(1 - \beta_0)$. **C.** Across simulations, feedback alignment in the cortex predicts learning performance (green circles), but feedback alignment in the thalamus is uncorrelated with learning performance (black circles). All models have N = 256 cortical neurons.

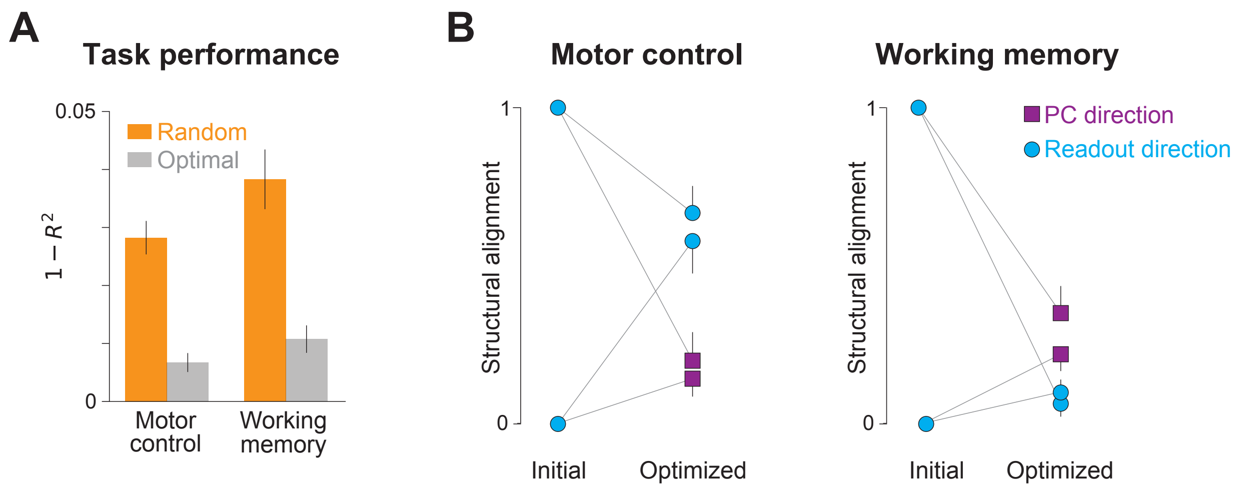


Figure S2: **Robustness of optimized corticothalamic weights determined by meta-learning**. **A.** Models with optimized corticothalamic connectivity outperform models with random connectivity even when both thalamocortical and readout weights are updated in parallel. In these models, the component of optimized corticothalamic weights aligned with readout weights is updated during the learning phase, while keeping the orthogonal components fixed. **B.** The degree of alignment of optimized corticothalamic weights with readout and principal component directions is robust to the alignment of the model at the beginning of meta-learning. In these models, corticothalamic weights are either orthogonal or perfectly aligned with the readout direction in the beginning of the meta-learning phase ("Initial") but the final alignments are qualitatively similar ("Optimized"). Error bars denote standard errors estimated by bootstrapping. All models have N = 256 cortical neurons and M = 32 thalamic neurons.

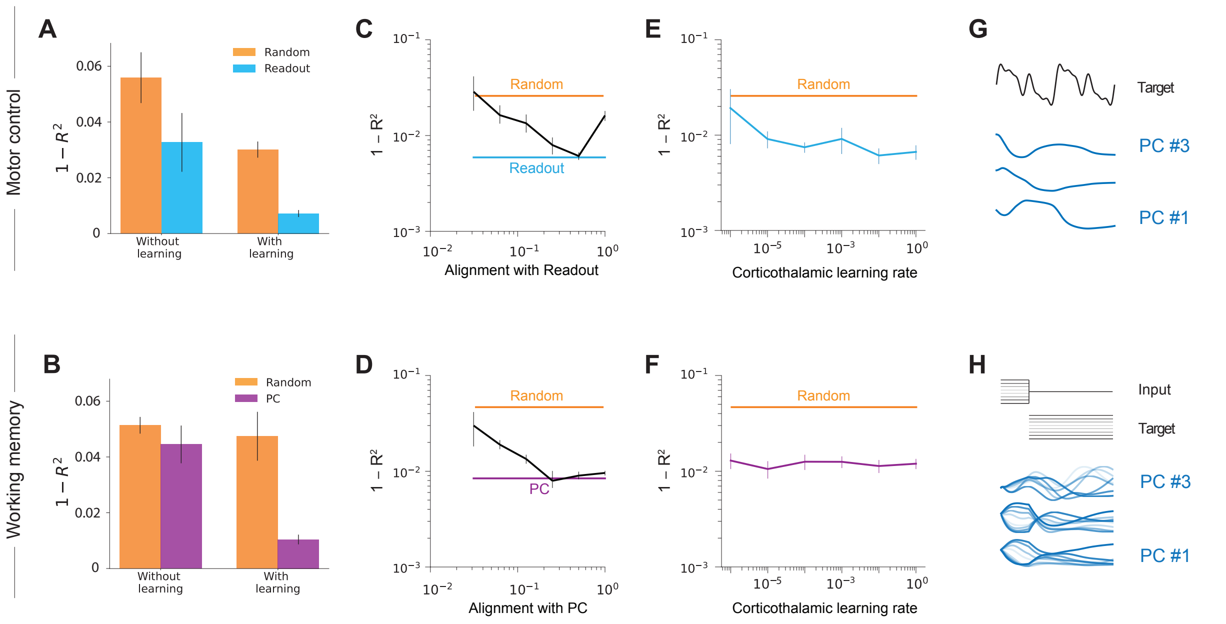


Figure S3: **Significance and robustness of subspace aligned corticothalamic connectivity models**. **A-B.** Task performance of models with and without learning in thalamocortical synapses. In the motor control task (**A**), models differed in terms of whether corticothalamic weights are random or aligned with readout, whereas in the working memory task (**B**), models differed in terms of whether corticothalamic weights are random or aligned with the principal component. Readout weights are updated in all models. **C-D.** Learning performance of models with partial alignment of corticothalamic weights with readout (**C**) or principal component directions (**D**). Different models were tested by systematically varying the degree of alignment of corticothalamic weights onto individual thalamic neuron with readout or principal component directions on a logarithmic scale. **C-D.** Learning performance of models in which corticothalamic weights were slowly updated to align with readout or principal component: $\mathbf{W}_k^{\mathrm{TC}} = (1-\beta) * \mathbf{W}_{k-1}^{\mathrm{TC}} + \beta * u$, where $u$ denotes either readout (**E**) or principal component (**F**), and $\beta$ denotes the speed of update. **G-H.** The target function (black), and the top three principal components of cortical activity (blue traces) at the beginning of learning. Individual traces in **H** correspond to response to different inputs. All models have N = 256 cortical neurons and M = 16 thalamic neurons.

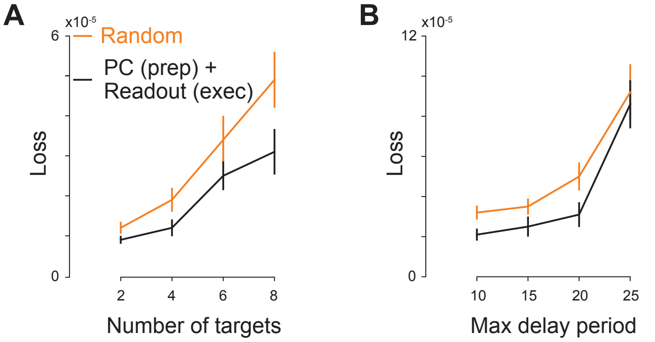


Figure S4: **Robustness of subspace aligned thalamocortical model of goal-directed reaching to task variables**. **A.** The models in which corticothalamic weights onto thalamic neurons active during preparation (execution) are aligned with the cortical principal component (readout) learn better than models with random corticothalamic weights, regardless of the number of reach targets. **B.** Similar to **A**, but showing learning performance as a function of the maximum delay between the location stimulus and the go cue. Note that in each condition, the delay was stochastic and drawn from a uniform distribution, $\mathcal{U}[5, t_{\max}]$, where $t_{\max}$ denotes the maximum delay period in units of neuronal time constant $\tau$.

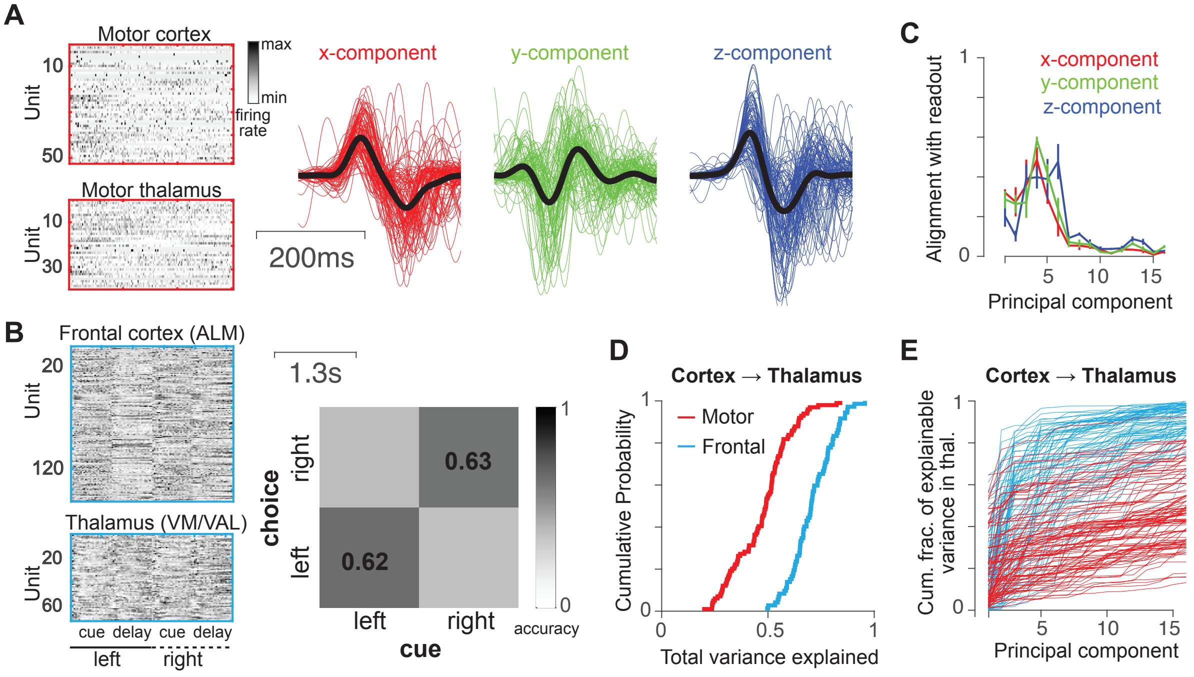


Figure S5: **Corticothalamic interactions in mice performing motor control and working memory tasks. A.** Left: Data from (Sauerbrei et al., 2020). Trial-averaged response matrix of neurons in the motor cortex (top) and motor thalamus (bottom) during the pellet grasping task, between onset and end of movement. Rows denote units and columns denote time. Response of each neurons is z-scored for the purpose of visualization. Right: Three spatial components of hand acceleration. Traces from individual trials are shown in color, and black traces denote the average across trials. **B.** Left: Data from (Guo et al., 2017). Similar to **A** (left), but during the delayed discrimination task. Response from trials with leftward and rightward responses were separately averaged and concatenated. Each trial comprised a ∼1.3s cue period followed by a ∼1.3s delay period. Right: Confusion matrix showing the choice accuracy in the leftward and rightward conditions. **C.** Mean alignment (across sessions) between the direction of readout weights and different cortical PCs, in the motor control task. Alignment of readout weights corresponding to each of the three spatial components of acceleration are shown separately (only the acceleration component with the highest variance (z-component) was shown in Figure 5). Error bars denote ±1 SEM. **D.** Cumulative probability distribution of the variance explained in the activity of thalamic neurons when decoding all neurons recorded from the cortex. Note that in all analysis, the ALM cortical population (n=151) was subsampled to match the dataset from motor cortex (n=53 ± 9). **E.** The fraction of explainable variance in individual thalamic neurons, as a function of the number of cortical principal components. Each curve denotes one thalamic neuron recorded during the motor control (red) and working memory (blue) tasks.
